# Accumulation of TCR signaling from self-antigens in naive CD8 T cells mitigates early responsiveness

**DOI:** 10.1101/2023.01.27.525946

**Authors:** Joel Eggert, Wendy M. Zinzow-Kramer, Yuesong Hu, Yuan-Li Tsai, Arthur Weiss, Khalid Salaita, Christopher D. Scharer, Byron B. Au-Yeung

**Affiliations:** Division of Immunology, Lowance Center for Human Immunology, Department of Medicine, Emory University; Department of Chemistry, Emory University; Rosalind Russell and Ephraim P. Engleman Rheumatology Research Center, Departments of Medicine and of Microbiology and Immunology, University of California, San Francisco; Department of Microbiology and Immunology, Emory University

**Author notes:** Correspondence to Byron B. Au-Yeung.

## Abstract

The cumulative effects of T cell receptor (TCR) signal transduction over extended periods of time influences T cell biology, such as the positive selection of immature thymocytes or the proliferative responses of naive T cells. Naive T cells experience recurrent TCR signaling in response to self-antigens in the steady state. However, how these signals influence the responsiveness of naive CD8^+^ T cells to subsequent agonist TCR stimulation remains incompletely understood. We investigated how naive CD8^+^ T cells that experienced relatively low or high levels of TCR signaling in response to self-antigens respond to stimulation with foreign antigens. A transcriptional reporter of *Nr4a1* (Nur77-GFP) revealed substantial heterogeneity of the amount of TCR signaling naive CD8^+^ T cells accumulate in the steady state. Nur77-GFP^HI^ cells exhibited diminished T cell activation and secretion of IFNγ and IL-2 relative to Nur77-GFP^LO^ cells in response to agonist TCR stimulation. Differential gene expression analyses revealed upregulation of genes associated with acutely stimulated T cells in Nur77-GFP^HI^ cells but also increased expression of negative regulators such as the phosphatase Sts1. Responsiveness of Nur77-GFP^HI^ cells to TCR stimulation was partially restored at the level of IFNγ secretion by deficiency of Sts1 or the ubiquitin ligase Cbl-b. Our data suggest that extensive accumulation of TCR signaling during steady state conditions induces a recalibration of the responsiveness of naive CD8^+^ T cells through gene expression changes and negative regulation, at least in part, dependent on Sts1 and Cbl-b. This cell-intrinsic negative feedback loop may allow the immune system to limit the autoreactive potential of highly self-reactive naive CD8^+^ T cells.

## Introduction

The activation of T cell-mediated immune responses is associated with sustained, robust signal transduction triggered by the T cell antigen receptor (TCR) (Courtney et al., 2018). Experienced over time, the cumulative effects of sustained TCR signaling build toward apparent signal thresholds required to cross essential checkpoints in the activation of T cell responses, including the commitment to enter a proliferative response (Allison et al., 2016; Clark et al., 2011; Preston et al., 2015). Naive T cells also experience TCR signals in secondary lymphoid organs (SLOs) in response to self-pMHC (Dorfman et al., 2000). These tonic or basal TCR signals are not associated with T cell activation but are experienced by naive CD4^+^ and CD8^+^ T cells constitutively in the steady state (This et al., 2022). How the cumulative effects of relatively weak or strong tonic TCR signals are interpreted by naive T cells and influence their responsiveness to subsequent foreign antigen stimulation remains unresolved (Myers et al., 2017b).

Tonic TCR signaling by naive T cells in response to self-pMHC is sufficient to induce constitutive tyrosine phosphorylation of the TCR complex and association of the tyrosine kinase Zap-70 with the CD3 ζ-chain (Stefanova et al., 2002; van Oers et al., 1994). Triggering of tonic TCR signals does not result in a cellular phenotype typically associated with an effector T cell (Myers et al., 2017b). However, tonic TCR signals can influence the expression of several genes at the transcriptional or protein level in T cells, including the cell surface molecules CD5 and Ly6C, and *Nr4a1*, which encodes the orphan nuclear receptor Nur77 (Mandl et al., 2013; Martin et al., 2013; Myers et al., 2017a). These findings suggest that the accumulation of varying levels of TCR signaling in naive T cells in the steady state can influence changes in T cell gene expression. This feature of tonic TCR signaling also raises the possibility that variable gene expression patterns in response to tonic TCR signaling result in functional heterogeneity within the naive T cell population (Eggert and Au-Yeung, 2021; Richard, 2022). This concept is consistent with models proposing that T cell responsiveness depends on previously experienced TCR signals (Huseby and Teixeiro, 2022). Taken to an extreme, relatively strong baseline TCR signaling could effectively result in T cell desensitization and hypo-responsiveness to subsequent TCR stimulations. Adaptive tuning in this context thus proposedly attenuates the responsiveness of the T cells within the naive T cell repertoire that respond most intensely to self-pMHC (Grossman and Paul, 1992).

Fluorescence-based reporters of *Nr4a* family genes, including *Nr4a1* (encoding Nur77) and *Nr4a3* (encoding Nor1), can provide fluorescence-based readouts of recently experienced TCR signaling (Jennings et al., 2020). The Nur77-GFP reporter transgene consists of enhanced green fluorescent protein (GFP) driven by the promoter and enhancer elements of the *Nr4a1* gene (Moran et al., 2011; Zikherman et al., 2012). A key feature of Nur77-GFP reporter expression is that GFP fluorescence intensity can reflect relative differences in TCR signal strength. For example, the mean fluorescence intensity of Nur77-GFP expressed by acutely stimulated T cells decreases with diminishing pMHC affinity (Au-Yeung et al., 2017; Moran et al., 2011). Furthermore, Nur77-GFP expression is relatively insensitive to constitutively active STAT5 or inflammatory signals, suggesting that reporter transgene expression is activated selectively by TCR stimulation in T cells (Moran et al., 2011). Moreover, TCR-induced Nur77-GFP expression depends on the function of intracellular mediators of TCR signaling, including the tyrosine kinase Zap-70. Previous work showed that stimulation with a single concentration of TCR stimulus in the presence of graded concentrations of a pharmacologic inhibitor of Zap-70 catalytic activity resulted in dose-dependent decreases in Nur77-GFP fluorescence intensity (Au-Yeung et al., 2014b).

Whereas some readouts of TCR signal transduction indicate signal intensity at a single time point, Nur77-GFP expression can reflect a relative level of TCR signal accumulation. For example, during thymic positive selection, CD4^+^ CD8^+^ double positive (DP) thymocytes experience multiple transient TCR stimulations over hours to days. DP thymocytes undergoing positive selection exhibit progressive increases in the level of Nur77-GFP, suggestive of a cumulative effect of multiple discrete TCR signaling events observed by transient calcium increases (Ross et al., 2014). Naive T cells express Nur77-GFP in the steady state, and in the CD4^+^ population, maintenance of Nur77-GFP expression depends on continuous exposure to MHCII (Moran et al., 2011; Zinzow-Kramer et al., 2019). In light of these findings, we propose that naive CD8^+^ T cells experience and adapt to the cumulative effects of tonic TCR signals.

In this study, we investigated the effects of accumulated TCR signaling on the functional responsiveness of naive CD8^+^ T cells. Naive CD8^+^ T cells expressing the highest levels of Nur77-GFP exhibit relative hypo-responsiveness to stimulation with agonist TCR ligands. Increased basal Nur77-GFP expression correlated with attenuated TCR-induced calcium fluxes, exertion of mechanical forces, and cytokine secretion compared with responses by Nur77-GFP^LO^ cells. Increases in accumulated TCR signaling were also associated with differential gene expression, including genes with the potential to inhibit T cell activation. We found that Nur77-GFP^HI^ cells from mice lacking *Ubash3b* (encoding Sts1) or Cbl-b exhibit partially rescued responsiveness to TCR stimulation. Together, these findings suggest a model whereby naive CD8^+^ T cells adapt to high levels of cumulative TCR signaling through negative regulation that limits initial T cell responsiveness.

## Results

### The accumulative TCR signaling from self-antigen in naive CD8^+^ T cells is heterogeneous

We first sought to investigate how diverse the accumulation of self-pMHC-driven TCR signaling is in the naive, CD44^LO^ CD62L^HI^ CD8^+^ T cell population. The distributions of Nur77-GFP fluorescence intensity of TCR polyclonal naive CD8^+^ and CD4^+^ T cells span over three orders of magnitude, as detected by flow cytometry (**Fig. 1 A**). By comparison, the GFP intensities of naive CD4^+^ and CD8^+^ T cells are notably higher than non-transgenic T cells but decreased compared to CD4^+^ Foxp3^+^ regulatory T cells (**Fig. 1 A**), a T cell population that expresses TCRs with high reactivity to self-pMHC (Hinterberger et al., 2010; Jordan et al., 2001; Lee et al., 2012). Moreover, GFP expression in naive CD8^+^ T cells positively correlates with the staining intensity of CD5 surface levels, a marker interpreted to correlate with TCR reactivity to self-pMHC (**Fig. 1 B**) (Cho et al., 2016; Mandl et al., 2013). These data suggest that naive CD4^+^ and CD8^+^ T cells accumulate varying amounts of TCR signaling in the steady state.

**Figure 1.**
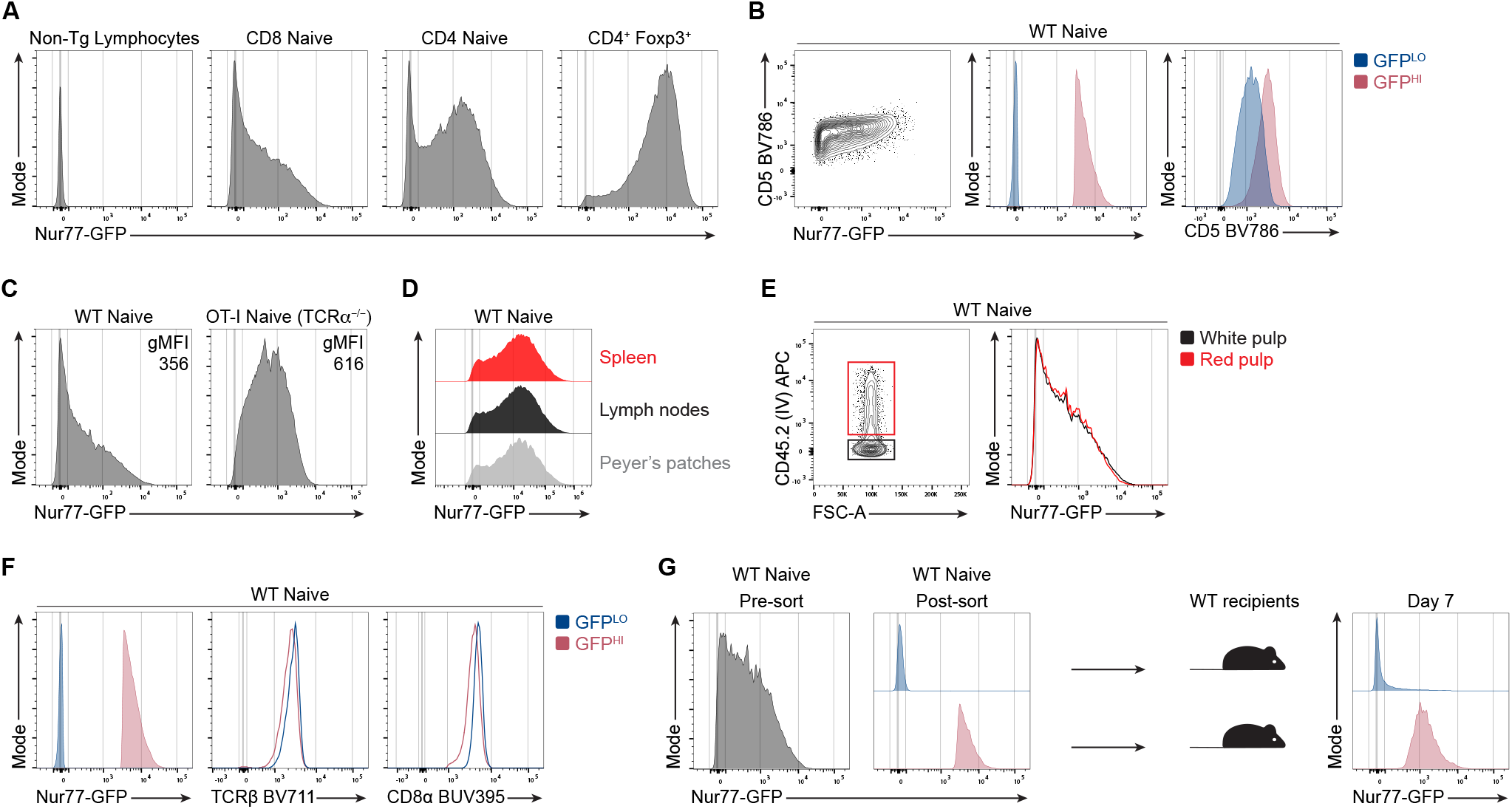
Accumulative TCR signaling in naive CD8^+^ T cells is heterogenous during steady-state conditions. **(A)** Representative flow cytometry plots of Nur77-GFP fluorescence of naive CD44^LO^ CD62L^HI^ CD8^+^ and CD4^+^ cells or CD4^+^ Foxp3-IRES-RFP^+^ T cells. All plots shown are from non-TCR transgenic mice **(B)** Contour plot (left) shows CD5 and Nur77-GFP expression by total naive polyclonal CD8^+^ T cells. Overlaid histogram (center) depicts GFP fluorescence for GFP^LO^ and GFP^HI^ cells. GFP^LO^ cells are the 10% of cells with the lowest (blue) GFP fluorescence intensity, whereas GFP^HI^ cells are the 10% of cells with the highest (red) GFP fluorescence intensity. Histogram (right) shows the CD5 expression for GFP^LO^ and GFP^HI^ populations. **(C)** Histograms show Nur77-GFP fluorescence intensities of naive CD8^+^ T cells from WT Nur77-GFP or OT-I-Nur77-GFP-TCRα^-/-^ mice. The numbers indicate the geometric mean fluorescence intensity (gMFI) calculated for the whole population. **(D)** Offset histograms show Nur77-GFP expression in naive polyclonal CD8^+^ T cells harvested from the spleen, mesenteric lymph nodes, or Peyer’s Patches. **(E)** Flow cytometry plots of naive polyclonal CD8^+^ T cells after intravascular labeling of cells in the red pulp by intravenous injection of CD45.2-APC antibody intravenously prior to euthanasia. **(F)** Histograms show expression of TCRβ and CD8α by polyclonal naive GFP^LO^ and GFP^HI^ CD8^+^ T cells. **(G)** Histograms show the GFP fluorescence intensity of total CD8^+^ T cells (left) or FACS-sorted GFP^LO^ and GFP^HI^ cells (middle). A total of 5×10^5^ GFP^LO^ or GFP^HI^ (top and bottom 10%) polyclonal CD8^+^ T cells were adoptively transferred into separate WT congenic recipients. Histogram (right) shows GFP fluorescence of transferred T cells seven days post-transfer and gated on naive CD8^+^ T cells and the congenic marker expression. Data represent two (**A, C, D, G**) to three (**B, E, F**) independent experiments with *n* = 2-3 mice.

We hypothesized that restricting the repertoire to a single TCR specificity would decrease the heterogeneity of GFP expression in a TCR transgenic population. To test the influence of TCR specificity on the distribution of GFP expression, we compared the intensity and distribution of GFP between naive polyclonal and OT-I TCRα^-/-^ TCR transgenic populations. The geometric mean fluorescence intensity (gMFI) of GFP expressed by naive CD44^LO^ CD62L^HI^ OT-I cells was higher than the GFP gMFI for polyclonal naive CD8^+^ cells (**Fig. 1 C**; and **Fig. S1**). However, OT-I and polyclonal naive CD8^+^ T cells exhibited a similar range of Nur77-GFP fluorescence intensity that spans over three orders of magnitude. These results suggest that TCR specificity can influence the intensity of TCR signaling experienced by individual T cells. However, the strength of TCR signaling in the steady state remains heterogeneous in a population that expresses identical TCRs.

We next asked whether GFP expression by naive CD8^+^ T cells varied between cells harvested from different anatomical locations. Hence, we analyzed naive CD8^+^ T cells from different secondary lymphoid organs (SLOs), such as the spleen, mesenteric lymph nodes, and Peyer’s patches, and compared the expression of GFP between these populations. However, we did not detect differences in the intensity or distribution of GFP expression (**Fig. 1 D**). Subsequently, we questioned whether the location within the spleen could still contribute to heterogenous Nur77-GFP expression in naive CD8^+^ T cells. To compare the GFP distribution of T cells located in the more vascularized red pulp versus the white pulp of the spleen, we performed intravascular labeling with fluorescently labeled anti-CD45 antibodies 3 minutes prior to euthanasia. We detected largely overlapping GFP intensities for naive, polyclonal CD8^+^ T cells labeled with anti-CD45 and cells not labeled with anti-CD45, interpreted to represent cells located in the red and white pulp, respectively (**Fig. 1 E)**. These results suggest that GFP^LO^ and GFP^HI^ cells are not skewed in their distribution between the red or white pulp in the spleen or the SLOs we analyzed.

We hypothesized that naive, CD8^+^ GFP^HI^ T cells accumulate more TCR signals due to increased surface levels of the TCR or the CD8 co-receptor. However, the 10% highest GFP-expressing cells expressed largely overlapping or slightly lower surface levels of the TCR β-chain and the CD8α co-receptor than the 10% lowest GFP-expressing cells (**Fig. 1 F**). Hence, increased GFP expression in naive CD8^+^ T cells does not positively correlate with increased surface TCR and CD8 levels.

We next questioned whether GFP^LO^ and GFP^HI^ T cells would maintain skewed intensities of GFP expression over several days. To test the stability of GFP expression, we sorted the 10% lowest and highest GFP-expressing naive polyclonal CD8^+^ T cells and adoptively transferred each population into congenic WT recipients (**Fig. 1 G**). One week post-transfer, GFP^LO^ and GFP^HI^ naive donor T cells sustained biased GFP expression. While some downregulation of GFP was present in the GFP^HI^ population, few GFP^LO^ cells upregulated GFP, suggesting that most GFP^LO^ cells tend to maintain low GFP expression (**Fig. 1 G**).

### Naive CD8^+^ T cells that experience extensive TCR signaling from self-antigen are hypo-responsive to TCR stimulation

To analyze the functional responsiveness of naive T cells that have accumulated varying amounts of TCR signaling from endogenous interactions, we isolated three populations across the GFP distribution (GFP^LO^, GFP^MED^, and GFP^HI^) from naive, polyclonal CD8^+^ T cells (**Fig. 2 A**). After 24 hours of stimulation with soluble anti-CD3 antibodies and splenocyte APCs, we performed an IFNγ-secretion assay. Approximately 25% of GFP^LO^ cells secreted IFNγ, whereas two-fold fewer GFP^MED^ and less than 1% of GFP^HI^ cells secreted IFNγ (**Fig. 2 B and C**). Hence, there was an apparent inverse correlation between the intensity of steady-state GFP expression and the magnitude of anti-CD3 induced IFNγ-secretion.

**Figure 2.**
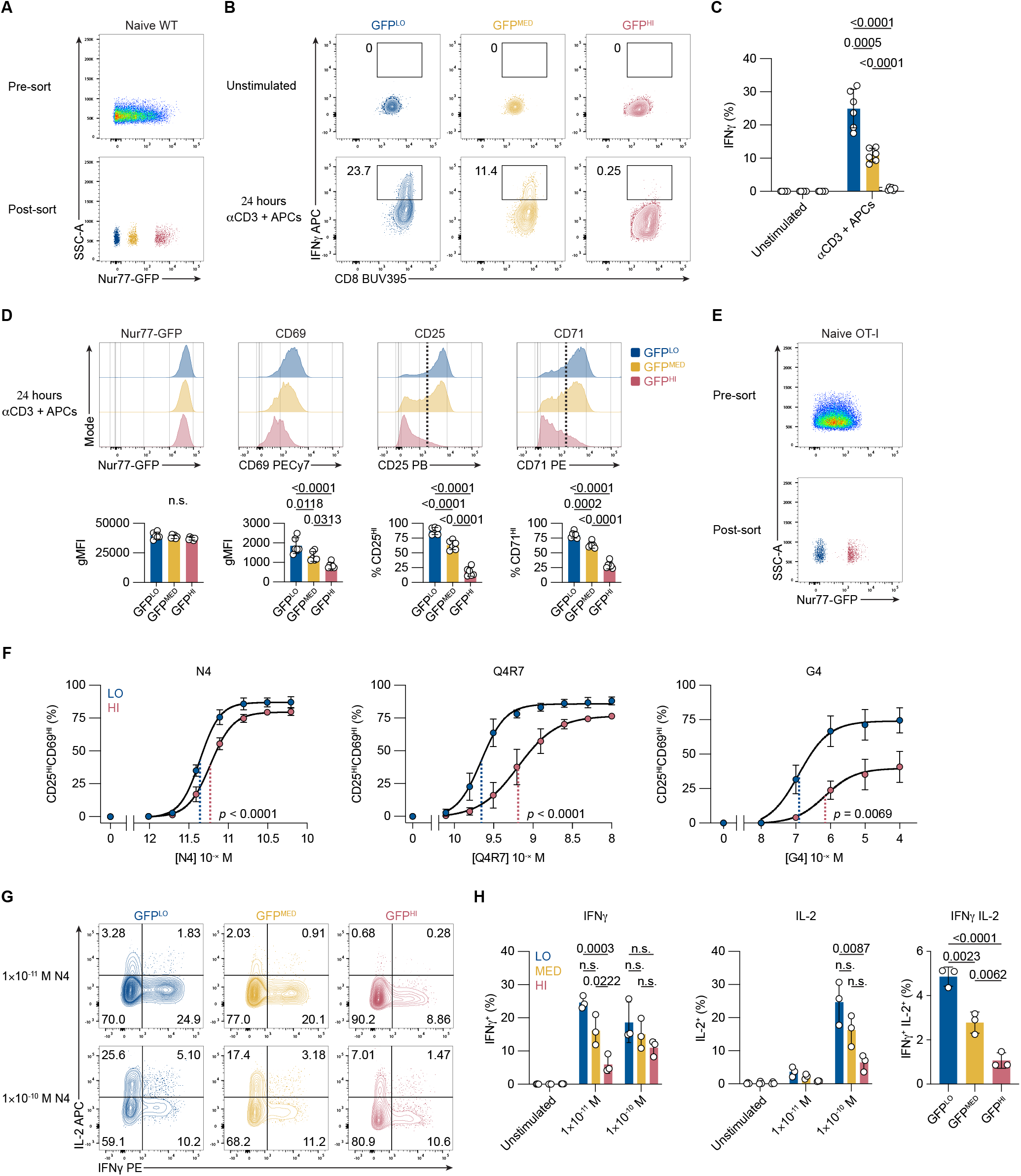
Accumulative steady-state TCR signaling correlates negatively with naive CD8 T cell responsiveness. **(A)** Representative flow cytometry plots show GFP fluorescence of total CD8^+^ cells (top) and sorted GFP^LO^, GFP^MED^, and GFP^HI^ naive, polyclonal CD8 T cell populations (bottom). **(B)** Contour plots depict CD8 and IFNγ expression by unstimulated and stimulated viable polyclonal CD8^+^ T cells after a 45 min IFNγ-secretion assay. Numbers indicate the percentage of cells within the indicated gates. **(C)** Bar graph displays the frequencies of GFP^LO^, GFP^MED^, and GFP^HI^ IFNγ-secreting cells. Cells were either unstimulated or stimulated for 24 hours with 0.25 μg/ml anti-CD3 and APCs before the secretion assay. Values are shown from three independent experiments. **(D)** Histograms show expression of the indicated activation markers of cells stimulated for 24 hours with 0.25 μg/ml anti-CD3. Cells were gated on viable CD8^+^ T cells. Bar graphs display the gMFI for Nur77-GFP and CD69 or the frequency of marker-positive cells for CD25 and CD71 (as indicated by the dotted line in the histogram). **(E)** Representative flow cytometry plots show GFP fluorescence for total naive OT-I CD8^+^ T cells pre-sorting (top), and FACS-sorted GFP^LO^ and GFP^HI^ cells (bottom). **(F)** Graphs show the frequencies of CD25^HI^CD69^HI^ cells after 16 hours of stimulation with indicated peptide concentrations and APCs. Plotted are mean values from three independent experiments fitted by non-linear regression curves. The dotted lines indicate the Log_10_ half maximal effective concentration (EC_50_) for GFP^LO^ (blue) and GFP^HI^ (red) cells. The *p*-value indicates the *t* test for the Log_10_EC_50_ (the null hypothesis being that the Log_10_EC_50_ is the same for the two populations). **(G)** Contour plots depict viable CD8^+^ T cells after a 45-minute assay of IFNγ- and IL-2-secretion of unstimulated or stimulated (16 hours) OT-I CD8^+^ T cells. **(H)** Bar graphs show the frequencies of IFNγ, IL-2, or IFNγ and IL-2-secreting cells after 16 hours of stimulation with indicated N4 peptide concentrations and APCs or unstimulated control. Values are shown from three independent experiments. All data represent three independent experiments with *n* = 3 mice (**E, F, G, H**) or *n* = 6 mice (**A, B, C, D**). Bars in **C**, **D**, and **H** depict the mean, and error bars show ± s.d. Statistical testing in **C** was performed by one-way analysis of variance (ANOVA) (p < 0.0001) followed by Tukey’s multiple comparisons test indicated in the graph. Statistical testing in **D** was performed by ANOVA (p < 0.0001 for CD69, CD25, and CD71) followed by Tukey’s multiple comparisons test. Statistical testing in **H** was performed by unpaired two-tailed Student’s *t* test. n.s., not significant.

To determine whether GFP^LO^, GFP^MED^, and GFP^HI^ cells similarly upregulated markers associated with acute T cell activation, we analyzed their expression of the activation markers CD25, CD69, and transferrin receptor (CD71), in addition to the Nur77-GFP reporter. All three populations upregulated Nur77-GFP and CD69 above baseline levels (**Fig. 2 D**; and **Fig S2 A**). However, on average, GFP^LO^ cells expressed higher levels of CD69 than GFP^MED^ and GFP^HI^ cells (**Fig. 2 D**). Similarly, higher frequencies of the GFP^LO^ population fully upregulated CD25 and CD71 (**Fig. 2 D**). Following stimulation, the sorted GFP^LO^, GFP^MED^, and GFP^HI^ populations each expressed similar levels of Nur77-GFP at the 24-hour endpoint. Considering their differential starting GFP MFIs, these results suggest that GFP^LO^ cells had experienced the highest cumulative amount of anti-CD3-induced TCR signaling compared to the GFP^MED^ and GFP^HI^ populations. Hence, the responsiveness to agonist TCR stimulation positively correlates with the net increase in Nur77-GFP expression from basal to endpoint level.

To test whether GFP^LO^ and GFP^HI^ cells exhibit differences in survival after stimulation, we quantified the proportion of viable CD8^+^ T cells after the 24-hour stimulation period. GFP^HI^ cells had a 1.5-fold reduction in the percentage of viable cells compared with GFP^LO^ cells (**Fig. S2 B**). Hence, GFP^HI^ cells experience a reduction in cell survival following TCR stimulation.

We next compared the effects of accumulated TCR:self-pMHC signaling on the responsiveness of naive CD8^+^ OT-I TCR transgenic cells with titrated doses of peptide and with altered peptides that vary in affinity for the OT-I TCR. We postulated that GFP^HI^ T cells exhibited decreased responsiveness for pMHC at low concentrations or weak affinity pMHC ligands. We applied the OT-I TCR transgenic system to test this hypothesis, utilizing the cognate SIINFEKL (N4) peptide and altered peptides with decreased affinities (Daniels et al., 2006). To compare GFP^LO^ and GFP^HI^ T cells expressing identical TCRs, we crossed OT1-Nur77-GFP mice to mice homozygous for the knockout allele of the endogenous TCR α-chain to prevent endogenous TCR recombination. Furthermore, we excluded Qa2^LO^ recent thymic emigrants (RTEs), which were more abundant in 6-9 week old OT-I-Nur77-GFP TCR transgenic mice, but present at only low frequencies in WT mice (**Fig. S2 C**). RTEs continue to undergo maturation and exhibit diminished functional responses compared to mature T cells (Boursalian et al., 2004). Thus, we sorted naive T cells with a CD8^+^ CD44^LO^ CD62L^HI^ Qa2^HI^ phenotype from OT-I TCR transgenic mice to compare mature T cell populations differing only in basal GFP expression. From this naive T cell population, we isolated the 10% lowest and highest GFP-expressing cells (**Fig. 2 E**). We assessed the upregulation of CD25 and CD69 after stimulating GFP^LO^ and GFP^HI^ OT-I cells for 16 hours with APCs and the cognate N4 peptide. The dose-response curve of GFP^HI^ cells was shifted further to the right compared to GFP^LO^ cells, indicating a reduction in CD25 and CD69 upregulation. The calculated Log_10_ EC_50_ value for GFP^LO^ cells was −11.36 compared to −11.23 for GFP^HI^ cells (**Fig. 2 F**; and **Fig. S2 D and E**). These results suggest that GFP^HI^ cells exhibit reduced responsiveness to a high-affinity antigen under non-saturating antigen doses.

To test whether the accumulation of extensive TCR signaling from self-pMHC affected the responsiveness to antigen affinity, we also stimulated OT-I cells with the SIIQFERL (Q4R7) altered peptide, which has reduced affinity for the OT-I TCR relative to the N4 peptide (Daniels et al., 2006). The dose-response curve of GFP^HI^ compared to GFP^LO^ cells was increasingly shifted to the right when stimulated with Q4R7 relative to N4. The calculated Log_10_ EC_50_ value for GFP^LO^ cells was −9.657 compared to −9.190 for GFP^HI^ cells (**Fig. 2 F**; and **Fig. S2 E**). Upon stimulation with the weak agonist peptide SIIGFEKL (G4), the dose-response curve also shifted to the right for GFP^HI^ cells. The calculated Log_10_ EC_50_ value for GFP^LO^ cells was −6.907 compared to −6.155 for GFP^HI^ cells (**Fig. 2 F**; and **Fig. S2 E**). These results indicate that higher levels of accumulated TCR signaling from self-pMHC in naive CD8^+^ T cells result in hypo-responsiveness to subsequent stimulation.

We next asked whether GFP^LO^ and GFP^HI^ cells exhibit differences in TCR-induced cytokine secretion. We hypothesized that GFP^HI^ cells would exhibit decreased IL-2 and IFNγ secretion relative to GFP^MED^ and GFP^LO^ cells. After sorting GFP^LO^, GFP^MED^, and GFP^HI^ OT-I cells and stimulating them for 16 hours with a concentration (1×10^-11^ M) of N4 peptide that was on the linear range of the curve for CD25- and CD69-upregulation, we performed IL-2- and IFNγ-capture assays (**Fig. 2 G and H**; and **Fig. S2 F and G**). GFP^LO^ OT-I cells generated the highest percentage of IFNγ-secreting cells (approximately 25%) (**Fig. 2 G and H**). There was a trend toward reduced IFNγ-secreting cells in the GFP^MED^ population (about 15%) and a significant reduction in the GFP^HI^ population (about 6%) (**Fig. 2 G and H**). The frequency of IL-2-secreting cells was below 5% for all populations at a dose of 1×10^-11^ M N4 peptide (**Fig. 2 G and H**).

To induce more robust IL-2 secretion, we stimulated the three populations with a ten-fold higher dose of N4 peptide (1×10^-10^ M). At this dose, there was comparable IFNγ secretion (**Fig. 2 G and H**). However, approximately 25% of GFP^LO^ cells secreted IL-2, whereas about 6% of GFP^HI^ cells secreted IL-2 (**Fig. 2 G and H**). Similarly, the frequency of cells that secreted both IL-2 and IFNγ was significantly higher in GFP^LO^ cells (about 5%) than in GFP^MED^ (approximately 2.5%) or GFP^HI^ cells (about 1%) (**Fig. 2 G and H**). Hence, there is a dose-dependent, inverse correlation between GFP expression in naive CD8^+^ T cells and cytokine secretion in response to subsequent foreign antigen stimulation.

We next asked whether GFP^LO^ and GFP^HI^ cells exhibit differences in cell division. We hypothesized that more accumulated TCR signaling from self-pMHC in naive CD8^+^ T cells would result in delayed or reduced cell division upon stimulation. We thus labeled CD8^+^ T cells with a cell proliferation dye and sorted naive GFP^LO^ and GFP^HI^ polyclonal T cells for *in vitro* stimulation with anti-CD3 antibodies and APCs (**Fig. S2 H and I**). Three days post-stimulation, the proliferation index (the average number of divisions of cells that divided at least once) of GFP^LO^ cells was greater than that of GFP^HI^ cells (**Fig. S2 I**). Together, these data suggest that the accumulation of TCR signaling from self-pMHC interactions negatively impacts the proliferative responses of naive CD8^+^ T cells under the conditions tested.

### Naive CD8^+^ GFP^LO^ and GFP^HI^ cells exhibit attenuated calcium flux responses and exert reduced mechanical forces

We next wanted to investigate whether GFP^HI^ cells exhibited an attenuated response at more proximal events of T cell activation upon stimulation with cognate peptide. Among the early T cell responses to pMHC stimulation is the exertion of mechanical forces through the TCR (Al-Aghbar et al., 2022). Previous work found a positive correlation between increases in the exertion of mechanical tension by T cells and increases in the intensity of Zap-70 phosphorylation, suggesting a positive regulatory role for mechanical forces in early T cell activation (Liu et al., 2016). We hypothesized that GFP^LO^ and GFP^HI^ cells would exhibit differences in tension exerted on pMHC ligands. To test this hypothesis, we utilized DNA hairpin-based “tension” probes linked to pMHC. The tension probe consists of a DNA hairpin conjugated to fluorophore (Atto647N) and quencher (BHQ2) molecules positioned to quench fluorescence by fluorescence resonance energy transfer (FRET) when the DNA hairpin is in its closed configuration (**Fig. 3 A**) (Ma et al., 2019). When a T cell, through its TCR, applies forces to a pMHC molecule with a magnitude exceeding 4.7 piconewtons (pN), the DNA hairpin unfolds, leading to the separation of the FRET pair and dequenching of the dye. A “locking” DNA strand is then introduced to selectively hybridize to the mechanically unfolded DNA hairpin and prevent refolding to capture the tension signal. After isolating the 10% lowest and highest GFP-expressing OT-I cells, we cultured them on substrates coated with tension probes conjugated to H2-K^b^ loaded with OVA N4 peptide. GFP^LO^ cells induced, on average, a 20% higher fluorescence signal from the tension probes than GFP^HI^ cells (**Fig. 3 B and C**). These results indicate that GFP^LO^ cells were more likely to exert the 4.7 pN tension force required to unfold the DNA hairpins than GFP^HI^ cells in response to pMHC stimulation.

**Figure 3.**
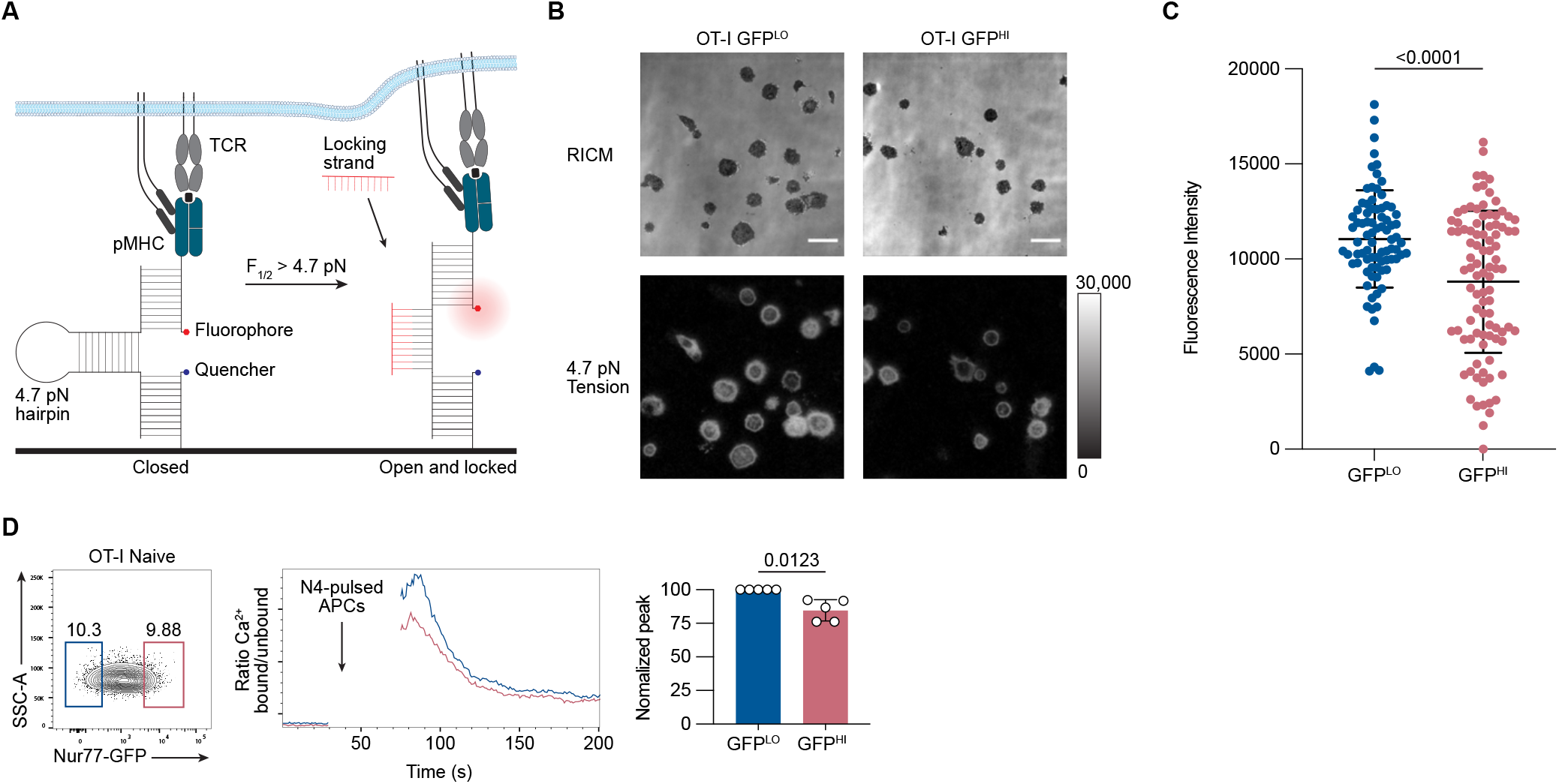
Nur77-GFP^HI^ CD8^+^ T cells exert less TCR-mediated tension forces and exhibit attenuated proximal TCR signaling. **(A)** Schematic outline of the DNA hairpin-based tension probe. In its closed conformation, the fluorescence of Cy3B is quenched. The DNA hairpin unfolds when TCR-mediated tension exceeds 4.7 piconewtons (pN). A “locking” DNA strand that hybridizes to the mechanically unfolded probe stabilizes the unfolded conformation of the DNA hairpin. **(B)** Representative Reflection Interference Contrast Microscopy (RICM) and fluorescence images showing GFP^LO^ and GFP^HI^ (top and bottom 10%) OT-I CD8^+^ T cells spread on DNA hairpin tension probe coated surfaces after 30 minutes. **(C)** Graph displays the unquenched fluorescence intensities of the unfolded tension probes for 81-94 cells. Each dot represents one cell. **(D)** Contour plot shows the distribution of Nur77-GFP fluorescence intensity for CD8^+^ CD44^LO^ OT-I T cells. Numbers indicate the percentages of cells within the indicated gates, representing GFP^LO^ and GFP^HI^ cells (left). Histogram shows the relative concentration of free Ca^2+^ over time. Shown are the mean values for GFP^LO^ and GFP^HI^ naive OT-I CD8^+^ T cells (middle). Baseline Ca^2+^ levels were recorded for 30 seconds, and the arrow indicates the time point when the T cells were mixed with N4-pulsed APCs, centrifuged, and resuspended before the continuation of data acquisition. The bar graph shows the normalized peak intracellular free Ca^2+^ values during ten seconds of GFP^LO^ and GFP^HI^ cells ~70 seconds after the initial acquisition (right). Data represent two **(C)** to three **(D)** independent experiments with *n* = 2 mice **(C)** or *n* = 5 mice **(D)**. Bars in **C** and **D** depict the mean, and error bars show ± s.d. Statistical testing was performed by unpaired two-tailed Student’s t test with Welch’s correction.

We next sought to determine whether GFP^LO^ and GFP^HI^ naive CD8^+^ T cells exhibit differences in proximal TCR signaling. We hypothesized that naive GFP^HI^ OT-I T cells would exhibit decreased cytosolic Ca^2+^ concentrations relative to GFP^LO^ cells upon stimulation with cognate N4 peptide antigen. Hence, we co-incubated OT-I cells labeled with the Indo-1 ratiometric indicator dye with N4 peptide-pulsed APCs and analyzed the fluorescent signal of the calcium indicator dye in T cells by flow cytometry. Compared to the peak free Ca^2+^ concentration signal generated by GFP^LO^ cells, the peak signal generated by GFP^HI^ cells was reduced by 20% (**Fig. 3 D**). Together, these data suggest that GFP^HI^ naive CD8^+^ T cells, which previously experienced more cumulative TCR signaling in the basal state, trigger downstream signals with weaker intensity in response to subsequent TCR stimulation. These results are consistent with a previous study using CD5 as a surrogate marker of self-pMHC reactivity, which showed an inverse correlation between the intensity of CD5 expression and the magnitude of anti-CD3-induced Ca^2+^ increases in naive CD8^+^ T cells (Cho et al., 2016).

### Extensive accumulation of TCR signaling in naive CD8 T cells correlates with differences in gene expression

To identify gene expression patterns associated with greater accumulation of TCR signaling in naive CD8^+^ T cells, we performed RNA-sequencing of naive CD8^+^ CD44^LO^ CD62L^HI^ Qa2^HI^ OT-I cells isolated based on the 10% highest versus 10% lowest GFP fluorescence intensities. We detected a total of 601 differentially expressed genes (DEGs) at a false discovery rate (FDR) < 0.05 (**Fig. 4 A**). Considering the correlation between Nur77-GFP expression and TCR signal strength, we hypothesized that GFP^HI^ cells would exhibit a gene expression profile with more similarities to acutely stimulated cells than GFP^LO^ cells. To test this hypothesis, we performed Gene Set Enrichment Analysis (GSEA) to compare our dataset of GFP^LO^ and GFP^HI^ naive CD8^+^ T cells with DEGs upregulated in viral infection-induced effector OT-I cells compared to naive cells (Luckey et al., 2006). Consistent with this hypothesis, GFP^HI^ cells showed enrichment of genes upregulated in effector CD8^+^ T cells (**Fig. 4 B**).

**Figure 4.**
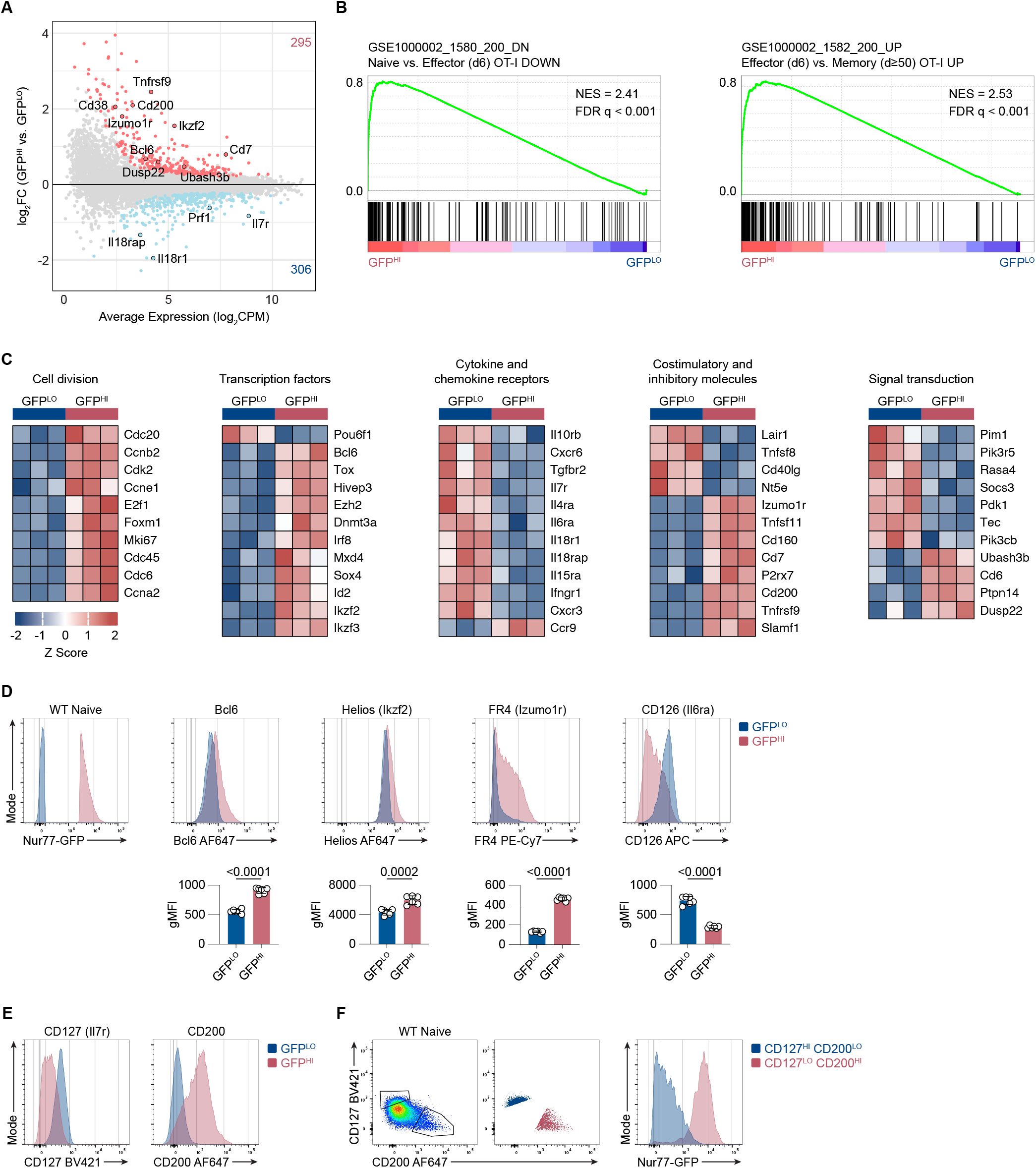
Nur77-GFP expression in naive CD8^+^ T cells during steady-state conditions correlates with gene expression changes. **(A)** MA plot of DEGs between GFP^LO^ and GFP^HI^ naive OT-I CD8^+^ T cells. DEGs were defined as genes with an FDR < 0.05. Selected genes have been highlighted. The number of upregulated and downregulated genes in GFP^HI^ relative to GFP^LO^ cells are indicated in red and blue, respectively. **(B)** GSEA of genes downregulated in naive compared to effector CD8^+^ T cells (left panel) and genes upregulated in effector compared to resting memory CD8^+^ T cells (right panel) (Luckey et al., 2006). FDR values were derived from running GSEA on the c7_Immunesigdb.v2022.1 database. **(C)** Curated heatmaps of normalized expression of DEGs in indicated categories. **(D)** Histograms show expression of the indicated markers by GFP^LO^ and GFP^HI^ cells. The cells were gated on naive, polyclonal CD8^+^ T cells. Bar graphs depict gMFI of indicated proteins. **(E)** Flow cytometry plots of CD127 and CD200 expression in naive GFP^LO^ and GFP^HI^ naive, polyclonal CD8^+^ T cells as in D. **(F)** Flow cytometry plots (left, middle) show the gating scheme to identify CD127^HI^ CD200^LO^ and CD127^LO^ CD200^HI^ populations. Histogram (right) shows the GFP fluorescence intensity for CD127^HI^ CD200^LO^ and CD127^LO^ CD200^HI^ populations. Plots depict naive, polyclonal Nur77-GFP CD8^+^ T cells. Data represent two (**D**) to three (**E** and **F**) independent experiments with *n* = 6 mice (**D**) or *n* = 3 mice (**E** and **F**). Bars in **D** depict the mean, and error bars show ± s.d. Statistical testing was performed by unpaired two-tailed Student’s t test. NES, normalized enrichment score.

Additionally, we compared the degree of overlap between DEGs in naive GFP^HI^ versus GFP^LO^ cells and DEGs in *Listeria* infection-induced OT-I effector cells versus naive OT-I cells (Best et al., 2013). Linear regression analysis indicated a significant correlation between genes enriched in GFP^HI^ cells and acutely stimulated OT-I cells (**Fig. S3 A**). These results suggest that accumulated TCR signaling from self-pMHC interactions in naive CD8^+^ T cells upregulates genes associated with acutely stimulated and effector CD8^+^ T cells. However, GFP^HI^ cells also showed enrichment of genes upregulated in effector compared to resting memory OT-I cells (**Fig. 4 B**). Therefore, recently experienced TCR stimulation may be a driver of the transcriptional differences between GFP^HI^ and GFP^LO^ naive CD8^+^ T cells.

We next sought to explore the sets of DEGs in GFP^HI^ naive CD4^+^ and CD8^+^ T cells. Therefore, we compared the DEGs between GFP^LO^ and GFP^HI^ naive CD8^+^ T cells and the DEGs upregulated in naive GFP^HI^ Ly6C^-^ CD4^+^ T cells (Zinzow-Kramer et al., 2022). Among the overlapping DEGs from both analyses (CD8^+^ and CD4^+^ cells), linear regression analysis suggested a significant correlation (**Fig. S3 B**). Hence, accumulating extensive TCR signals during steady-state conditions induces similar transcriptional changes in naive CD4^+^ and CD8^+^ T cells.

In addition, we detected increased transcripts of genes involved in cell division in GFP^HI^ relative to GFP^LO^ cells, consistent with a gene signature indicative of acutely activated T cells (**Fig. 4 C**). In agreement, naive CD8^+^ T cells that experience stronger tonic TCR signals and express higher levels of CD5 also show enrichment for cell cycle-associated genes (White et al., 2016). GFP^HI^ cells also expressed higher levels of transcription factors associated with T cell differentiation, such as *Bcl6* and *Ikzf2* (Helios), and TCR stimulation, such as *Tox* and *Irf8* (**Fig. 4 C**) (Alfei et al., 2019; Kaech and Cui, 2012; Miyagawa et al., 2012). Consistent with a gene signature of T cell activation, GFP^HI^ cells upregulated immunomodulatory molecules such as *Tnfrsf9* (4-1bb), *Tnfsf11* (Rankl), and *Cd200* (**Fig. 4 C**) (Pollok et al., 1995; Pollok et al., 1993; Snelgrove et al., 2008; Wong et al., 1997). GFP^HI^ cells expressed lower levels of *Il7r* (CD127) in addition to other common γ-chain cytokine receptors such as *Il4ra, Il6ra* (CD126), and *Il15ra* (**Fig. 4 C**). Among genes involved in signal transduction, GFP^HI^ cells had lower expression levels of kinases such as Pim1 and Pdk1. In contrast, GFP^HI^ cells expressed higher levels of the phosphatases *Ubash3b* (*Sts1*), *Dusp22* (Jkap), and *Ptpn14* (**Fig. 4 C**). Taken together, gene expression patterns associated with higher levels of accumulated TCR signaling bear similarities to gene expression patterns induced by acute TCR stimulation. This gene signature includes higher expression levels of immunomodulatory receptors, ligands, and negative regulators of TCR signaling.

We next performed flow cytometry analyses to determine whether differential gene expression patterns correlated with differential protein expression. We analyzed the 10% highest vs. lowest GFP-expressing naive, polyclonal CD8^+^ T cells to compare the expression of several DEGs, including *Bcl6, Ikzf2* (Helios), *Izumo1r* (Folate receptor 4), *Il6ra* (CD126), *Il7ra* (CD127), and *Cd200* (**Fig. 4 D and E**; and **Fig. S3 C**). For five of the six selected DEGs, protein staining was increased in GFP^HI^ relative to GFP^LO^ cells and thus correlated with the RNA-sequencing data. GFP^HI^ cells expressed lower surface levels of CD126, which was also consistent with the RNA-seq analysis. Flow cytometry analysis of naive CD8^+^ T cells showed a spectrum of CD127 and CD200 expression (**Fig. 4 E**). Within the naive CD8^+^ population, the CD127^HI^ CD200^LO^ cell subset enriched for Nur77-GFP^LO^ cells and in contrast, the CD127^LO^ CD200^HI^ population enriched for GFP^HI^ cells (**Fig. 4 F**). These results indicate that Nur77-GFP^LO^ and GFP^HI^ cells exhibit differential expression of several genes at the protein level.

### Sts1 negatively regulates the responsiveness of GFP^LO^ and GFP^HI^ naive CD8^+^ cells

Differential gene expression analyses revealed that Nur77-GFP^HI^ naive OT-I cells expressed higher levels of *Ubash3b* (encoding Sts1), a phosphatase that negatively regulates TCR signaling (Mikhailik et al., 2007). We hypothesized that the absence of Sts1 in GFP^HI^ naive CD8^+^ T cells would increase the responsiveness of GFP^HI^ cells. To test this hypothesis, we analyzed CD127^HI^ CD200^LO^ (GFP^LO^-like) and CD127^LO^ CD200^HI^ (GFP^HI^-like) naive CD8^+^ T cells isolated from WT and Sts1^-/-^ mice. Notably, the percentages of GFP^LO^-like and GFP^HI^-like cells were similar in WT and Sts1^-/-^ mice (**Fig. S4 A**). Isolated GFP^LO^-like and GFP^HI^-like CD44^LO^ CD62L^HI^ CD8^+^ T cells were stimulated with APCs and anti-CD3 antibodies for 24 hours and analyzed for upregulation of CD25, CD69, and IFNγ secretion (**Fig. S4 B**). Following stimulation, the frequency of cells that upregulated CD25 and CD69 was approximately 50% in the WT GFP^LO^-like population compared to approximately 7% in the WT GFP^HI^ population (**Fig. 5 A**). Hence, GFP^LO^-like cells were more responsive than GFP^HI^-like cells, which recapitulates the attenuated response of Nur77-GFP^HI^ cells compared to Nur77-GFP^LO^ cells (**Fig. 2**). In the absence of Sts1, the frequency of cells that upregulated CD25 and CD69 increased in both GFP^LO^-like and GFP^HI^-like populations (**Fig. 5 A**; and **Fig. S4 C**). The ratio of Sts1^-/-^ to WT cells that upregulated CD25 and CD69 was similar for both GFP^LO^-like and GFP^HI^-like populations (**Fig. 5 B**). These analyses suggest that Sts1 decreases the responsiveness of both GFP^LO^-like and GFP^HI^-like populations.

**Figure 5.**
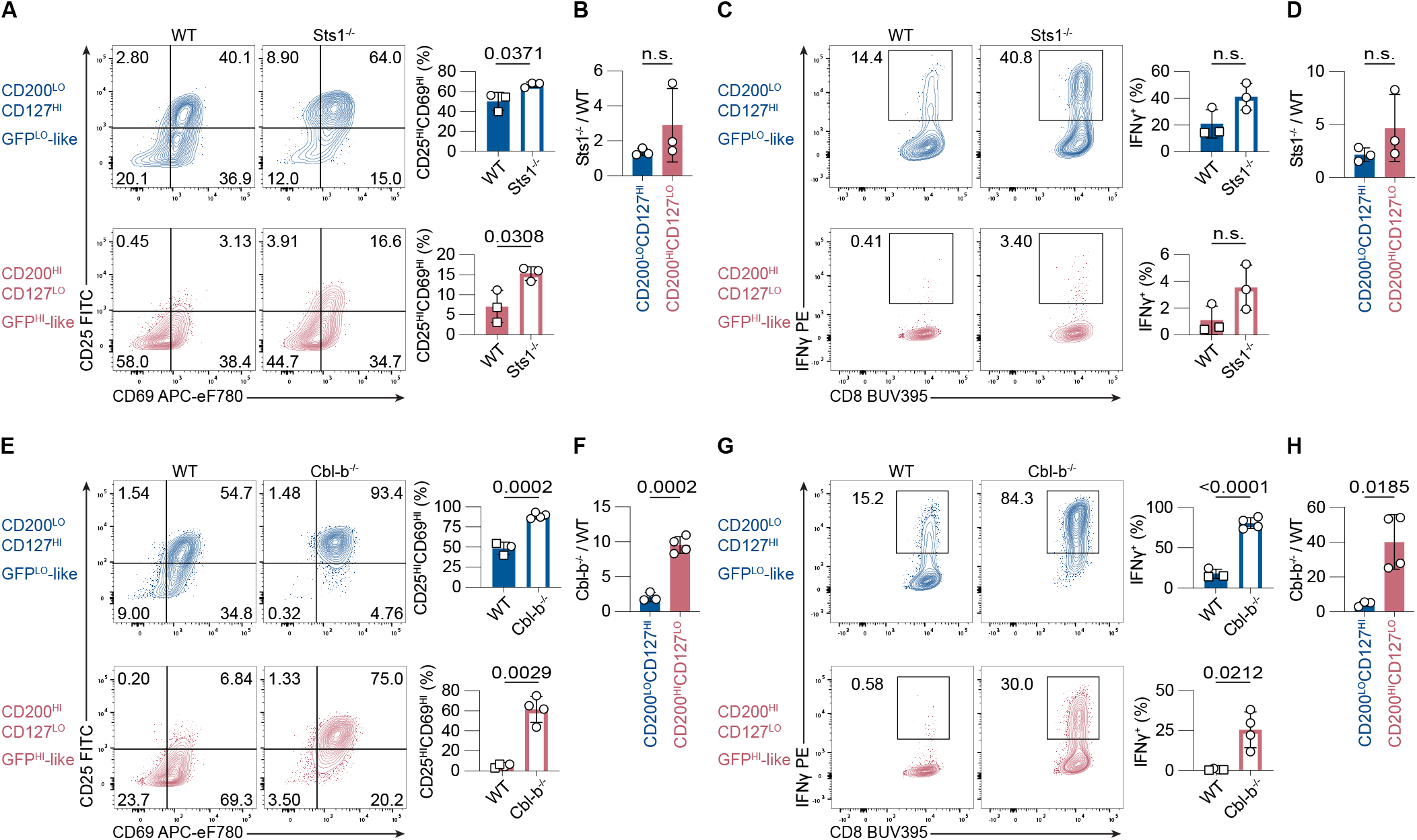
Sts1 and Cbl-b contribute to the attenuated responsiveness of naive Nur77-GFP CD8^+^ T cells. **(A** and **E)** Contour plots depict CD25 and CD69 upregulation in naive, polyclonal GFP^LO^-like and GFP^HI^-like CD8^+^ T cells stimulated for 24 hours with 0.25 μg/ml anti-CD3 and APCs. Numbers indicate the percentage of cells within each quadrant. Bar graphs depict the frequencies of CD25^HI^CD69^HI^ cells from three to four independent experiments. **(B** and **F)** Bar graphs show the ratio of %CD25^HI^CD69^HI^ Sts1^-/-^ cells (in **B**) or Cbl-b^-/-^ cells (in **F**) to %CD25^HI^CD69^HI^ WT cells, within GFP^LO^-like (blue) and GFP^HI^-like (red) populations. **(C** and **G)** Contour plots of IFNγ-secretion of CD8^+^ T cells stimulated as in **A** and **E**, after a 45-minute IFNγ-secretion assay. Numbers indicate the frequency of cells within the IFNγ^+^ gate. Bar graphs show the percentages of IFNγ^+^ cells from three to four independent experiments. **(D** and **H)** Bar graph shows the ratio of the frequencies of Sts1^-/-^ (in **D**) or Cbl-b^-/-^ (in **H**) versus WT IFNγ-secreting cells within the GFP^LO^-like (blue) and GFP^HI^-like (red) cell populations. Some of the WT data points for experiments in **A**, **C**, **E**, and **G** overlap since two experiments with Sts1^-/-^ and Cbl-b^-/-^ T cells were conducted simultaneously. Overlapping WT data points are labeled with squares instead of circles. Data represent three to four independent experiments with *n* = 3-4 mice. All bars depict the mean and error bars depict ± s.d. Statistical testing was performed by unpaired two-tailed Student’s *t* test in **A**, **C**, **E** (upper panel), **F**, and **G** (upper panel). Statistical testing was performed by unpaired two-tailed Student’s *t* test with Welch’s correction in **B**, **D**, **E** (lower panel), **G** (lower panel), and **H**. n.s., not significant.

To examine whether Sts1 deficiency could rescue IFNγ-secretion in GFP^HI^-like cells, we performed an IFNγ-capture assay after the 24-hour stimulation period (**Fig. 5 C**; and **Fig. S4 D**). There was a non-significant difference but a trend toward increased IFNγ-secreting cells in Sts1^-/-^ vs. WT GFP^LO^-like cells (**Fig. 5 C**). Similarly, there was a non-significant trend toward a higher frequency of IFNγ-secreting cells in GFP^HI^-like Sts1^-/-^ (about 3%) compared to WT cells (about 1 %) (**Fig 5 C**). The ratio of IFNγ-secreting cells between Sts1^-/-^ and WT CD8^+^ T cells was similar in GFP^LO^-like vs. GFP^HI^-like cells (**Fig. 5 D**). These data suggest that the phosphatase Sts1 limits the responsiveness of both GFP^LO^-like and GFP^HI^-like naive CD8^+^ T cells to some degree.

### Cbl-b deficiency partially rescues the responsiveness of GFP^HI^ naive CD8^+^ T cells

Previous mass spectrometry studies revealed that Sts1 associates with Cbl-b, an E3 ubiquitin ligase (Voisinne et al., 2016). Cbl-b is also a negative regulator of TCR signaling, and Cbl-b deficiency results in CD28-independent T cell activation and increased susceptibility to autoimmune diseases (Li et al., 2019). Considering the interaction between Sts1 and Cbl-b and the inhibitory function of Cbl-b in the TCR signal transduction pathway, we hypothesized that Cbl-b deficiency would rescue the attenuated responsiveness of GFP^HI^ cells. Compared to WT naive CD8^+^ T cells, naive Cbl-b^-/-^ T cells contain similar percentages of GFP^LO^-like and GFP^HI^-like cells (**Fig. S4 E**). For these studies, we also utilized CD127 and CD200 surface expression to isolate GFP^LO^-like and GFP^HI^-like naive CD8^+^ T cells, and we stimulated these populations for 24 hours with APCs and anti-CD3 antibodies (**Fig. S4 F**). In both GFP^LO^-like and GFP^HI^-like cell populations, the percentages of cells that fully upregulated CD25 and CD69 were higher in Cbl-b-deficient samples than in WT samples (**Fig. 5 E**; and **Fig. S4 G**). The frequency of CD25^HI^ CD69^HI^ cells was 1.5-fold higher in Cbl-b^-/-^ compared to WT GFP^LO^-like cells (**Fig. 5 E**). Likewise, while only 5% of WT GFP^HI^-like cells fully upregulated CD25 and CD69, the frequency was more than ten-fold higher in Cbl-b^-/-^ GFP^HI^-like cells. The ratio of CD25^HI^ CD69^HI^ cells between Cbl-b^-/-^ and WT CD8^+^ T cells was more than four-fold higher in GFP^HI^-like compared to GFP^LO^-like cells (**Fig. 5 F**). These data suggest that the responses of GFP^HI^-like cells were rescued to a greater extent by Cbl-b deficiency than GFP^LO^-like cells.

We next asked whether Cbl-b deficiency could also rescue the secretion of IFNγ in GFP^HI^-like cells. After 24 hours of stimulation with anti-CD3-mediated TCR-crosslinking, we performed an IFNγ-capture assay. We observed a higher frequency of IFNγ-secreting cells in Cbl-b^-/-^ compared to WT T cells (**Fig. 5G**; and **Fig. S4 H**). IFNγ secretion in GFP^LO^-like Cbl-b-deficient T cells was about four-fold more prevalent compared to GFP^LO^-like WT cells (**Fig. 5G**). Similarly, approximately 20% of GFP^HI^-like Cbl-b^-/-^ T cells secreted IFNγ, while the frequency of IFNγ-secreting cells was less than 1% in the GFP^HI^-like WT population (**Fig. 5G**). The ratio of IFNγ-secreting cells in Cbl-b^-/-^ compared to WT T cells was almost nine-fold higher in GFP^HI^-like vs. GFP^LO^-like cells (**Fig. 5H**). These results indicate that Cbl-b-deficiency results in hyperresponsiveness of all naive CD8^+^ T cells to TCR stimulation. However, Cbl-b deficiency appears to have a larger effect in rescuing the responsiveness of GFP^HI^ cells than GFP^LO^ cells. Together, these data support a model where the accumulation of extensive self-pMHC-induced TCR signals induces negative regulation, in part, mediated by Cbl-b.

## Discussion

In this study, we found that basal expression levels of a Nur77-GFP reporter transgene inversely correlate with the initial responsiveness of naive CD8^+^ T cells to stimulation with agonist TCR ligands. Higher levels of accumulated TCR signaling correlated with changes in gene expression, including the upregulation of genes that could negatively regulate signal transduction in T cells. Hence, we propose that naive CD8^+^ T cells that experience extensive TCR:self-pMHC signals over time induce negative feedback mechanisms that limit their responsiveness to subsequent TCR stimulations.

Previous studies of Nur77-GFP reporter expression during T cell development showed that following positive selection, DP and CD8^+^SP thymocytes express elevated levels of GFP compared to pre-selection thymocytes (Zikherman et al., 2012). Still, the distribution of GFP fluorescence intensity spanned over three orders of magnitude in polyclonal and OT-I TCR transgenic positively selected thymocytes (Au-Yeung et al., 2014a). These findings raised the possibility that a subset of T cells induce relatively low or high levels of Nur77-GFP expression during their development in the thymus. In this study, we detected a similarly broad distribution of GFP fluorescence intensity within the naive CD8^+^ T cell population. Together, these findings open the possibility that some T cells experience relatively weak or strong TCR signals constitutively, first as immature T cells in the thymus and then as naive T cells in the secondary lymphoid organs. Thus, some CD8 SP thymocytes that initially experience strong tonic TCR signaling during development may continue to experience strong tonic TCR signaling as naive CD8^+^ T cells in the steady state. We hypothesize that the GFP^HI^ population, to some degree, is comprised of the naive T cell population that experienced strong TCR signaling in the thymus but escaped negative selection. This is consistent with early studies showing rapid Nur77 expression during negative selection of thymocytes (Cheng et al., 1997). Recent studies have also provided evidence for mature CD8^+^ T cells that recognize self-pMHC but exhibit reduced functionality or tolerance (Truckenbrod et al., 2021). Such tolerogenic responses are evident in T cells that experience constitutive agonist TCR stimulation in mice unperturbed by infection or inflammatory mediators (Trefzer et al., 2021).

Recent studies suggest that *Nr4a* factors, including *Nr4a1* (encoding Nur77), have a role in restraining peripheral T cell responses (Odagiu et al., 2020). Consistent with this concept, in vivo-tolerized murine T cells express high levels of *Nr4a1, Nr4a1* overexpression results in the upregulation of anergy-associated genes such as Cbl-b whereas *Nr4a1* deficiency results in resistance to anergy induction and exacerbation of autoimmune disease severity (Hiwa et al., 2021; Liebmann et al., 2018; Liu et al., 2019). Moreover, *Nr4a1^-/-^ Nr4a2^-/-^ Nr4a3^-/-^* CAR T cells had an enhanced antitumor response in a solid tumor mouse model (Chen et al., 2019). These studies suggest that *Nr4a1* and the other *Nr4a* family genes can act as negative regulators. We propose that the transcriptional upregulation of *Nr4a1* in Nur77-GFP^HI^ naive CD8^+^ cells is indicative of negative feedback that attenuates their responsiveness.

Our differential gene expression analyses suggested that accumulation of strong tonic TCR signaling induced upregulation of genes associated with acute TCR stimulation, as well as the phosphatases *Ubash3b* (encoding Sts1), *Dusp22* (encoding Jkap), and *Ptpn14* which have the potential to function as negative regulators of intracellular signaling in naive OT-I GFP^HI^ cells. The phosphatase Jkap can dephosphorylate kinases of the proximal TCR signaling cascade, while Ptpn14 has unclear functions in T cells (Li et al., 2014; Stanford et al., 2012). We found that *Sts1* deficiency partially rescues the responsiveness of GFP^HI^ naive CD8^+^ T cells, consistent with previous work showing that *Sts1^-/-^* and *Sts1^-/-^ Sts2^-/-^* T cells are hyperresponsive to TCR stimulation (Carpino et al., 2004; Mikhailik et al., 2007). The modest effect of *Sts1* deficiency on cytokine production by GFP^HI^ cells suggested that there could be more dominant factors that drive their hyporesponsive phenotype. Sts1’s role in negatively regulating the responsiveness of GFP^HI^ T cells may involve the inhibition of Zap-70 through the dephosphorylation of regulatory tyrosine residues (Mikhailik et al., 2007).

In addition to the phosphatase function of Sts1, it is also possible that the ubiquitin ligase function of Cbl-b mediates the negative regulation of GFP^HI^ cells (Lutz-Nicoladoni et al., 2015). Another possibility is that the post-translational regulation and activity of Cbl-b differs between GFP^LO^ and GFP^HI^ cells. Moreover, since Sts1 is an interaction partner of Cbl-b, it is possible that Cbl-b could facilitate the recruitment of Sts1 to TCR-proximal tyrosine kinases (Voisinne et al., 2016). Further studies are required to investigate how the contribution of the phosphatase activity of Sts1, the ubiquitin ligase activity of Cbl-b, and the bridging function of the Sts1:Cbl-b interaction contribute to the attenuated responsiveness induced by cumulative TCR signaling in T cells.

Variable levels of Nur77-GFP expression appear to correlate with functional heterogeneity within the naive CD8^+^ T cell population. It is possible that negative regulation of naive T cells with increased reactivity to self-pMHC influences such variations at the single-cell level. Lineage tracing studies have previously identified diversity in the expansion and differentiation of single T cells through primary and recall responses (Buchholz et al., 2016). Cellular heterogeneity may also contribute to the dynamic nature of adaptive immune responses to respond to a breadth of antigens (Richard, 2022; Wong and Germain, 2018).

In conclusion, we observed reduced responsiveness in naive CD8^+^ T cells that accumulated high levels of TCR:self-pMHC stimulation in the steady state. Extensive TCR signaling mediated by self-antigen interactions promotes negative regulation dependent, at least in part, on the phosphatase Sts1 and the ubiquitin ligase Cbl-b. We speculate that such negative feedback mechanisms may constitute a form of cell-intrinsic tolerance in naive T cells.

## Materials and Methods

### Mice

Nur77-GFP (Tg(Nr4a1-EGFP)GY139Gsat) transgenic mice, Zap-70 deficient mice lacking mature T cells (Zap70tm1Weis), and Foxp3-RFP mice (C57BL/6-Foxp3tm1Flv/J) have been previously described (Kadlecek et al., 1998; Wan and Flavell, 2005; Zikherman et al., 2012). C57BL/6J mice (WT mice in the text) and CD45.1 mice (B6.SJL-Ptprca Pepcb/BoyJ) were purchased from the Jackson Laboratory. A Nur77-GFP strain that is interbred with the OT-I (C57BL/6-Tg(TcraTcrb)1100Mjb/J) TCR transgenic strain was described previously (Au-Yeung et al., 2017). This OT-I-Nur77-GFP strain was interbred with a TCRα^-/-^ strain (B6.129S2-Tcratm1Mom/J) purchased from the Jackson Laboratory. A Nur77-GFP strain interbred with the Foxp3-RFP strain has previously been described (Zinzow-Kramer et al., 2019). All mice were housed under specific pathogen-free conditions in the Division of Animal Resources at Emory University. Sts1^-/-^, and Cbl-b^-/-^ strains were described previously (Carpino et al., 2004; Chiang et al., 2000). These strains were maintained in the Laboratory Animal Resource Center at the University of California, San Francisco. Both female and male mice were used throughout the study. All animal experiments were conducted in compliance with the Institutional Animal Care and Use Committees at Emory University and the University of California, San Francisco.

### Antibodies and reagents

For negative enrichment of CD8^+^ T cells, the following biotinylated anti-mouse or anti-mouse/human antibodies were purchased from BioLegend: CD4 (clone RM4-5), CD19 (6D5), B220 (RA3-6B2), CD11b (M1/70), CD11c (N418), CD49b (DX5), and Erythroid cells (TER119). For negative selection of APCs, biotinylated anti-CD4 (RM4-5), CD8α (53-6.7), and Erythroid cells (TER119) were purchased from BioLegend. For flow cytometry, anti-CD126 (clone D7715A7), CD19 (6D5), CD25 (PC61), CD4 (RM4-5), CD44 (IM7), CD45.1 (A20), CD45.2 (104), CD62L (MEL-14), CD8 (53-6.7), Qa-2 (695H1-9-9), and TCR β chain (H57-597) were purchased from BioLegend. Anti-Bcl6 (clone K112-91), CD127 (SB/199), CD200 (OX-90), CD44 (IM7), CD5 (53-7.3), CD62L (MEL-14), CD71 (C2), CD8α (53-6.7), and Helios (22F6) were purchased from BD Biosciences. Anti-CD69 (clone H1.2F3) and FR4 (eBio12A5) were purchased from ThermoFisher Scientific. Streptavidin conjugated to APC (catalog #SA1005) and eFluor 450 (catalog #48-4317-82) were purchased from ThermoFisher Scientific. For viability, LIVE/DEAD fixable Near-IR, Violet or Yellow (ThermoFisher Scientific) was used according to the manufacturer’s instructions.

### Lymphocyte isolation and flow cytometry

Single-cell suspensions of lymphoid organs were generated by mashing organs through a 70 μm cell strainer or using a Dounce homogenizer. For phenotypic analysis of T cells by flow cytometry, red blood cells (RBCs) were lysed using RBC Lysis Buffer (Tonbo Biosciences) prior to Fc-block incubation (anti-mouse CD16/CD32, clone 2.4G2, Tonbo Biosciences). CD8^+^ T cells were purified by negative selection using biotinylated antibodies and magnetic beads, as previously described (Smith et al., 2016). Splenocytes were used as APCs, isolated from Zap70^-/-^ or TCRα^-/-^ mice after RBC lysis or by negative selection using biotinylated antibodies and magnetic beads on single-cell suspensions from C57BL/6 mice. Single-cell suspensions were stained in PBS and washed with FACS buffer (PBS with 0.5% BSA and 2 mM EDTA) for surface stains. For intracellular staining, samples were fixed and permeabilized with the Foxp3/Transcription Factor Staining kit according to the manufacturer’s instructions (ThermoFisher Scientific). For in vitro proliferation analysis, T cells were labeled with CellTrace Violet (ThermoFisher Scientific) according to the manufacturer’s instructions. Samples were analyzed using FACSymphony A5 (BD Biosciences), LSRFortessa (BD Biosciences), or Cytek Aurora instruments. Flow cytometry data were analyzed using FlowJo v.10.8.1 software (BD Biosciences).

### Intravascular labeling

Intravascular labeling was performed as previously described (Anderson et al., 2012). Briefly, 3 μg anti-CD45.2-APC antibody was injected in 200 μl PBS intravenously, 3 min prior to euthanasia. Cells from the spleen were analyzed by flow cytometry. Lymph nodes and peripheral blood were harvested as negative and positive controls, respectively. Positive staining with anti-CD45 antibodies was interpreted to indicate cells located within the red pulp; the absence of staining with anti-CD45 was interpreted to indicate cells located within the white pulp.

### Cell sorting

Naive CD8^+^ GFP^LO^ and GFP^HI^ T cells were sorted from bulk CD8^+^ T cells using a FACS Aria II SORP cell sorter (BD Bioscience). From viable polyclonal CD8^+^ CD44^LO^ CD62L^HI^ cells, the 10% of cells with the highest and the 10% of cells with the lowest GFP fluorescence intensity were sorted. For OT-I cells, samples were sorted on GFP expression (top and bottom 10%) from viable CD8^+^ CD44^LO^ CD62L^HI^ Qa2^HI^ cells. For the DNA hairpin tension probe experiment, bulk CD8^+^ T cells were sorted based on a viable, CD4^-^ CD19^-^ phenotype, then GFP^LO^ and GFP^HI^ cells were isolated from the 10% of cells with the highest and lowest GFP fluorescence intensity. The purity of CD8^+^ T cells post-enrichment was >96%.

### Adoptive transfer

For the Nur77-GFP stability experiment, 5×10^5^ sorted naive, polyclonal GFP^LO^ or GFP^HI^ CD8^+^ T cells were injected intravenously into congenic WT recipients in 200 μl PBS. Flow cytometry analysis was conducted seven days later on CD8^+^ T cells enriched from the spleen and lymph nodes.

### T cell stimulation

For *in vitro* stimulation of T cells, 5 × 10^4^ sorted CD8^+^ T cells were cultured with 2.5 × 10^5^ APCs (T cell-depleted splenocytes) in a 96-well U-bottom plate. Polyclonal CD8^+^ T cells were incubated with 0.25 μg/ml anti-CD3ε antibodies (clone, ID), whereas OT-I cells were incubated with SIINFEKL (N4) or SIIQFERL (Q4R7) or SIIGFEKL (G4) peptides (GeneScript) at indicated concentrations. Cells were cultured in RPMI 1640 (ThermoFisher Scientific) supplemented with 10% FBS, % L-Glutamine, % Pen/Strep, % HEPES, % Sodium Pyruvate, % non-essential Amino Acids, and % 2-mer-capto-ethanol at 37°C with 5% CO^2^.

### Cytokine secretion assay

IFNγ-secreting polyclonal CD8^+^ T cells were labeled using the IFNγ Secretion Assay Kit (Miltenyi Biotech, catalog #130-090-984) after 24 hours of stimulation with APCs and peptide. IFNγ- and IL-2-secreting OT-I cells were co-labeled using the IFNγ Secretion Assay Kit (Miltenyi Biotech, catalog #130-090-516) and the IL-2 Secretion Assay Kit (Miltenyi Biotech, catalog #130-090-987) after 16 hours of stimulation. Briefly, 1-1.5 × 10^5^ T cells, including co-cultured T cell-depleted splenocytes, were labeled with the bispecific catch reagent and incubated in 50 ml of pre-warmed RPMI supplemented with 10% FBS for 45 min at 37°C. 50 ml conical tubes were inverted every 5 minutes several times during incubation. After washing, cells were stained with the cytokine detection antibody/antibodies in addition to surface antibodies.

### Calcium analysis

OT-I cells were labeled with 1.5 μM Indo-1 AM dye (ThermoFisher Scientific) according to the manufacturer’s instructions. APCs (T cell-depleted splenocytes) were pulsed for 30 minutes at 37°C with 1 μM SIINFEKL peptide and washed. All cells were incubated at 37°C during the acquisition and for 5 min before the start of the experiment. After the baseline calcium levels of 4 × 10^6^ OT-I cells were recorded for 30 seconds, cells were pipetted to an Eppendorf tube containing 8 × 10^6^ peptide-pulsed APCs and spun down for 5 seconds in a microcentrifuge. The acquisition was resumed after the cell pellet was resuspended. The ratio of bound dye (Indo-violet) to unbound dye (Indo-blue) was analyzed for the 10% top and bottom GFP-expressing cells gated on viable CD8^+^ CD44^LO^ cells.

### Preparation of tension probe surfaces

No. 1.5H glass coverslips (Ibidi) were placed in a rack and sequentially sonicated in Milli-Q water (18.2 megohms cm-1) and ethanol for 10 minutes. The glass slides were then rinsed with Milli-Q water and immersed in freshly prepared piranha solution (3:1 sulfuric acid:H_2_O_2_) for 30 minutes. The cleaned substrates were rinsed with Milli-Q water at least six times in a 200-mL beaker and washed with ethanol thrice. Slides were then incubated with 3% 3-aminopropyltriethoxysilane (APTES) in 200 mL ethanol for 1 hour, after which the surfaces were washed with ethanol three times and baked in an oven at 100°C for 30 minutes. The slides were then mounted onto a six-channel microfluidic cell (Sticky-Slide VI 0.4, Ibidi). To each channel, ~50 mL of NHS-PEG4-azide (10 mg/ml) in 0.1 M NaHCO_3_ (pH 9) was added and incubated for 1 hour. Afterward, the channels were washed with 1 mL Milli-Q water three times, and the remaining water in the channel was removed by pipetting. The surfaces were then blocked with 0.1% BSA for 30 minutes and washed with PBS three times. Subsequently, the hairpin tension probes were assembled in 1 M NaCl by mixing the Cy3B-biotin labeled ligand strand (Atto647N - CGC ATC TGT GCG GTA TTT CAC TTT - Biotin) (220 nM), DBCO-BHQ2 labeled quencher strand (DBCO-TTT GCT GGG CTA CGT GGC GCT CTT – BHQ2) (220 nM), and hairpin strand (GTG AAA TAC CGC ACA GAT GCG TTT GTA TAA ATG TTT TTT TCA TTT ATA CTTTAA GAG CGC CAC GTA GCC CAG C) (200 nM) in the ratio of 1.1:1.1:1. The mixture was heat-annealed at 95°C for 5 minutes and cooled down to 25°C over a 30-minute time window. The assembled probe (~50 mL) was added to the channels (Final concentration = 100 nM) and incubated overnight at room temperature. This strategy allows for covalent immobilization of the tension probes on azide-modified substrates via strain-promoted cycloaddition reaction. Unbound DNA probes were washed away by PBS the next day. Then, streptavidin (10 mg/ml) was added to the channels and incubated for 45 minutes, followed by washes with PBS. Next, a biotinylated pMHC (OVA N4-H2k^b^) ligand (10 mg/ml) was added to the surfaces, incubated for 45 minutes, and washed with PBS. Surfaces were buffer exchanged with Hanks’ balanced salt solution before imaging.

### Imaging TCR tension with DNA hairpin tension probes

TCR:pMHC interactions exert force and mechanically unfold the DNA hairpin, leading to the dye’s (Atto647N-BHQ2) dequenching. T-cells were added to the tension probe surface and incubated for 20 minutes at room temperature. 200 nM of locking strand was then added to the surface for 10 minutes to capture the tension signal.

### RNA-Sequencing

1 × 10^5^ CD8^+^ CD44^LO^ CD62L^HI^ Qa2^HI^ OT-I GFP^LO^ and GFP^HI^ cells from three biological replicates were sorted into RLT Lysis Buffer (Qiagen) containing 1% 2-mercaptoethanol. RNA was isolated using the Zymo Quick-RNA MicroPrep kit (Zymo Research), cDNA was prepared from 1000 cell equivalent of RNA using the SMART-Seq v4 Ultra Low Input RNA Kit for Sequencing (Takara Bio), and next-generation sequencing libraries were generated using the Nextera XT DNA Library Preparation kit (Illumina). The library size patterning from a 2100 Bioanalyzer (Agilent) and the DNA concentration were used as quality control metrics of the generated libraries. Samples were sequenced at the Emory Nonhuman Primate Genomics Core on a NovaSeq6000 (Illumina) using PE100. FastQC (https://www.bioinformatics.babraham.ac.uk/projects/fastqc/) was used to validate the quality of sequencing reads. Adapter sequences were trimmed using Skewer, and reads were mapped to the mm10 genome using STAR (Dobin and Gingeras, 2015; Jiang et al., 2014). Duplicate reads were identified using PICARD (http://broadinstitute.github.io/picard/) and were removed from the following analyses. Reads mapping to exons were counted using the R package GenomicRanges (Lawrence et al., 2013). Genes were considered expressed if three reads per million were detected in all samples of at least one experimental group.

Analysis of differentially expressed genes was conducted in R v.4.1.1 using the edgeR package v.3.36.0 (Robinson et al., 2010). Genes were considered differentially expressed at a Benjamini-Hochberg FDR-corrected *p*-value < 0.05. Heatmaps were generated using the ComplexHeatmap v.2.10.0 R package (Gu et al., 2016). Preranked GSEA was conducted using the GSEA tool v.4.2.3 (Subramanian et al., 2005). The ranked list of all detected transcripts was generated by multiplying the sign of the fold change by the –Log_10_ of the *p*-value. All other RNA sequencing plots were generated using the ggplot2 v.3.3.5 R package (Wickham, 2016).

## Statistical analysis

All statistical analyzes were performed in Prism v.9.4.1 (GraphPad) or R v.4.1.1. A *p*-value < 0.05 was considered significant. Details about the statistical tests used is available in each figure legend. The sample sizes of experiments were determined based on preliminary experiments or prior experiments with CD4^+^ T cells that yielded significant results. No power analyzes to calculate sample sizes were performed.

## Data availability

RNA sequencing data are available under accession number GSE223457 in the Gene Expression Omnibus (https://www.ncbi.nlm.nih.gov/geo/query/acc.cgi?acc=GSE223457).

## Acknowledgements

We thank Simon Grassmann and Wan-Lin Lo for critical reading of the manuscript, the Pediatric/Winship Flow Cytometry Core for cell sorting and the Emory Integrated Genomics Core (EIGC) for RNA-sequencing. This work was supported in part by the National Institute of Allergy and Infectious Diseases R01AI165706 (to B.B.A-Y.), NCI T32 CA108462-17 (Y.-L.T.) and Cancer Research Institute Irvington Postdoctoral Fellowship (Y.-L.T.).

## Author contributions

J.E. and B.B.A.-Y. conceptualized the study. J.E., W.Z.-K., and Y.H. performed experiments. J.E., W.Z.-K., and C.D.S. analyzed the RNA-sequencing data. K.S. and Y.H. designed and performed the tension probe experiments. Y.-L.T. and A.W. contributed conceptual input and provided Sts1^-/-^ and Cbl-b^-/-^ cells. J.E. and B.B.A.-Y. wrote the manuscript with input from all authors.

### Disclosures

A.W. is a co-founder and a scientific advisory board member of Nurix Therapeutics, Inc., which has a Cbl-b inhibitor in phase 1 clinical trials. A.W. owns stock and receives consulting fees from Nurix. No other disclosures were reported.

---

Figure S1. **Representative gating of naive CD8^+^ T cells.** Representative gating of naive polyclonal and naive OT-I CD8^+^ T cells. Numbers indicate the percentage of cells within each gate.

Figure S2. **Accumulative steady-state TCR signaling correlates negatively with naive CD8 T cell responsiveness, supporting data. (A)** Histograms show expression of the indicated activation markers of unstimulated control cells. **(B)** The frequency of viable CD8^+^ T cells was determined after 24 hours of stimulation with 0.25 μg/ml anti-CD3 and APCs. **(C)** Contour plots depict Qa2 and CD8 expression in naive polyclonal or naive OT-I CD8^+^ T cells in mice aged 6-9 weeks. Numbers indicate the percentage of cells within the indicated gates. Bar graph shows the frequency of Qa2^LO^ cells in WT or OT-I mice. **(D)** Unstimulated control of CD25 and CD69 upregulation in GFP^LO^ and GFP^HI^ naive OT-I cells. **(E)** Representative flow cytometry plots depicting CD25 and CD69 upregulation after 16 hours of stimulation with indicated peptide concentrations are shown from one experiment. Panels in the first row represent suboptimal peptide concentrations, the second row depicts peptide concentrations on the linear part of the dose-response curve, and the third row show saturating peptide concentrations. Numbers indicate the percentage of cells within the indicated gates. **(F)** Representative flow cytometry plots of the GFP distribution of pre-sort, total OT-I T cells (left) and sorted GFP^LO^, GFP^MED^, and GFP^HI^ naive, OT-I CD8 T cell populations (right). **(G)** Sorted naive OT-I cells and APCs were incubated for 45 minutes as an unstimulated control for the IFNγ- and IL-2 secretion assay. **(H)** Representative flow cytometry plots of the pre-sort GFP distribution (left) and sorted GFP^LO^ and GFP^HI^ naive, polyclonal CD8 T cell populations (right). **(I)** CTV-labeled naive, polyclonal GFP^LO^, and GFP^HI^ CD8^+^ T cells were incubated for 70 hours with 0.25 μg/ml anti-CD3 and APCs. The representative flow cytometry plot was gated on viable CD8^+^ T cells. The graph depicts the proliferation index (the average number of divisions of cells that divided at least once) of four independent experiments. Data represents two (**C**), three (**A**, **D**, **E**, **F**, **G**), or four (**B**, **H**, **I)** independent experiments with *n* = 2 (**C**), *n* = 3 (**D**, **E**, **F**, **G**), *n* = 4 (**A**, **H**, **I**) or *n* = 6 (**A**) mice. Bars in **B**, **C**, and **I** depict the mean and error bars show ± s.d. Statistical testing was performed by unpaired twotailed Student’s t test in **B** and **I** or by unpaired two-tailed Student’s *t* test with Welch’s correction in **C**.

Figure S3. **Nur77-GFP^HI^ naive CD8^+^ T cells have transcriptional changes associated with T cell activation. (A)** Log_2_ fold-change plot of genes upregulated in effector compared to naive OT-I CD8^+^ T cells on the Y-axis (Best et al., 2013) and genes upregulated in Nur77-GFP^HI^ compared to GFP^LO^ naive OT-I CD8^+^ T cells on the X-axis. Each dot represents an overlapping DEG defined as genes with an FDR < 0.05 present in both datasets. The red line depicts the correlation with a 95% confidence interval. The dotted black line depicts a 1:1 relationship between the two datasets. **(B)** Similar to A, the plot depicts the Log_2_ fold-change of genes upregulated in Nur77-GFP^HI^ compared to GFP^LO^ naive Ly6C^-^ CD4^+^ T cells on the Y-axis (Zinzow-Kramer et al., 2022) and genes upregulated in GFP^HI^ compared to GFP^LO^ naive OT-I CD8^+^ T cells on the X-axis. **(C)** The top row depicts Nur77-GFP expression in relationship to indicated markers in naive, polyclonal CD8^+^ T cells. The bottom row indicates the Fluorescence Minus One (FMO) control for the indicated markers. Data are from two independent experiments from *n* = 6 mice. Statistical analysis in **A** and **B** was performed by a one-sample *t* test (the null hypothesis being that the slope was equal to zero).

Figure S4. **Sts1 and Cbl-b contribute to the attenuated responsiveness of naive Nur77-GFP CD8^+^ T cells, supporting data. (A)** The expression of CD127 and CD200 in naive, polyclonal CD8^+^ T cells from WT and Sts1^-/-^ mice. **(B)** Representative flow cytometry plots of sorted, naive GFP^LO^-like (blue) and GFP^HI^-like cells (red) CD8^+^ T cells from WT and Sts1^-/-^ mice. **(C)** CD25 and CD69 expression in unstimulated naive cells as indicated from WT or Sts^-/-^ mice. **(D)** Sorted naive, polyclonal CD8^+^ T cells and APCs were incubated for 45 minutes as an unstimulated control for the IFNγ-secretion assay. **(E)** The expression of CD127 and CD200 in naive, polyclonal CD8^+^ T cells from WT and Cbl-b^-/-^ mice. **(F)** Representative flow cytometry plots of sorted, naive GFP^LO^-like (blue) and GFP^HI^-like cells (red) CD8^+^ T cells from WT and Cbl-b^-/-^ mice. **(G)** CD25 and CD69 expression in unstimulated naive cells as indicated from WT or Cbl-b^-/-^ mice. **(H)** Sorted naive, polyclonal CD8^+^ T cells and APCs were incubated for 45 minutes as an unstimulated control for the IFNγ-secretion assay. All data represents 3-4 experiments with *n* = 3-4 mice.

