## Supplementary figures and images for "Accumulation of TCR signaling from self-antigens in naive CD8 T cells mitigates early responsiveness"

### Supplemental Figures

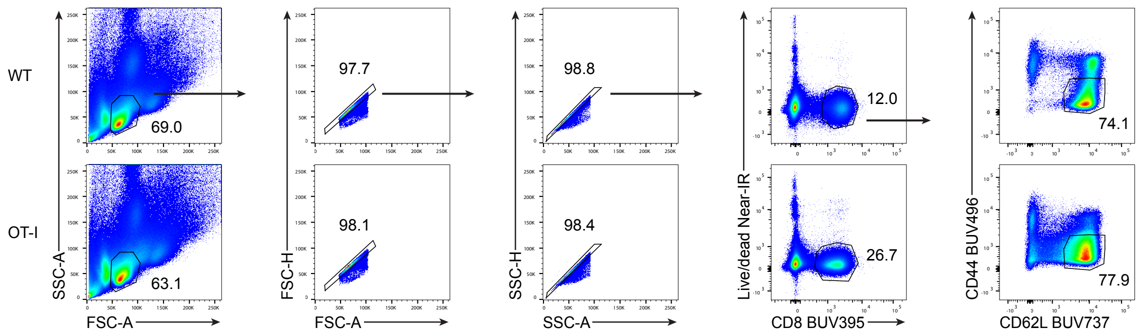

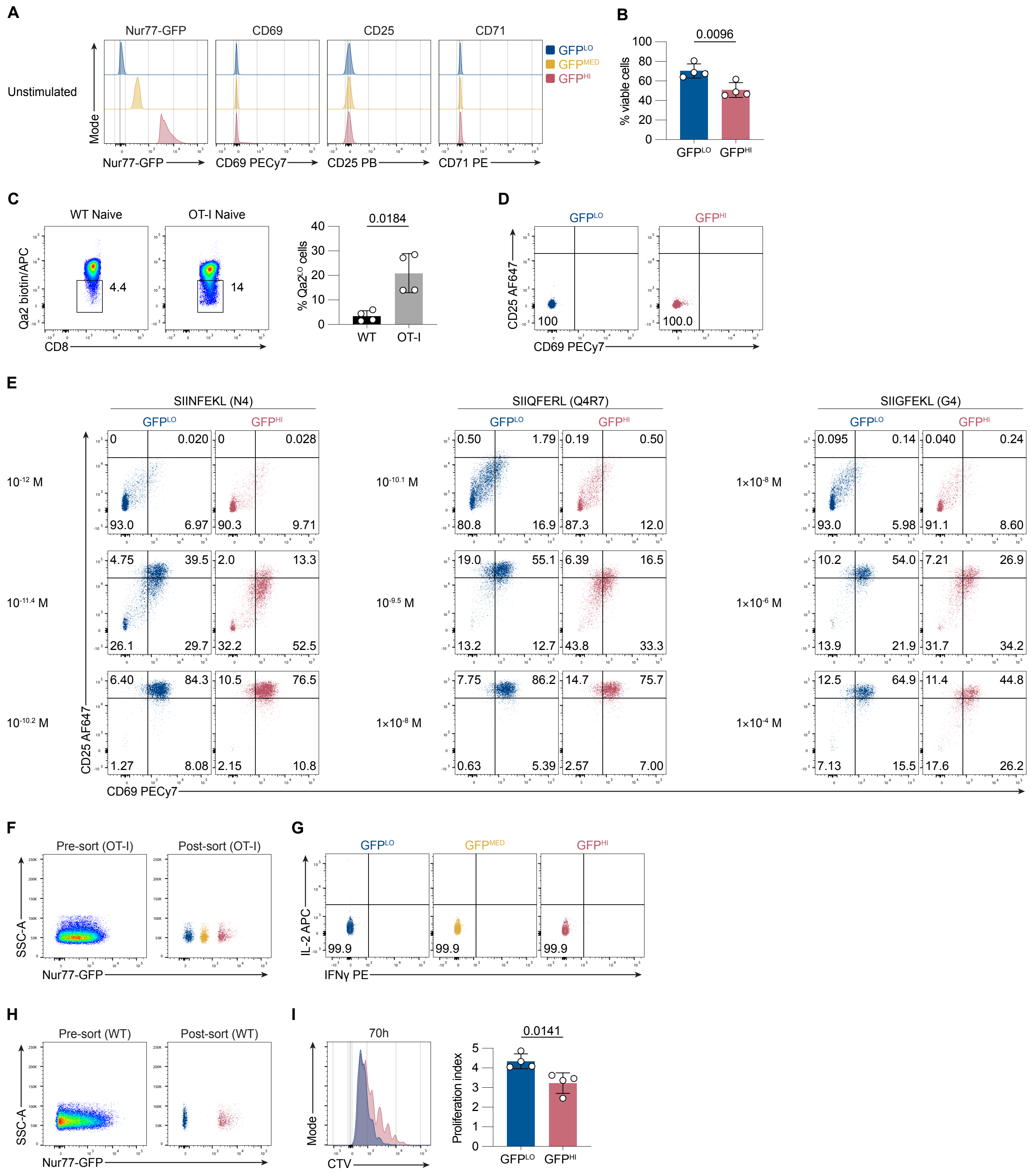

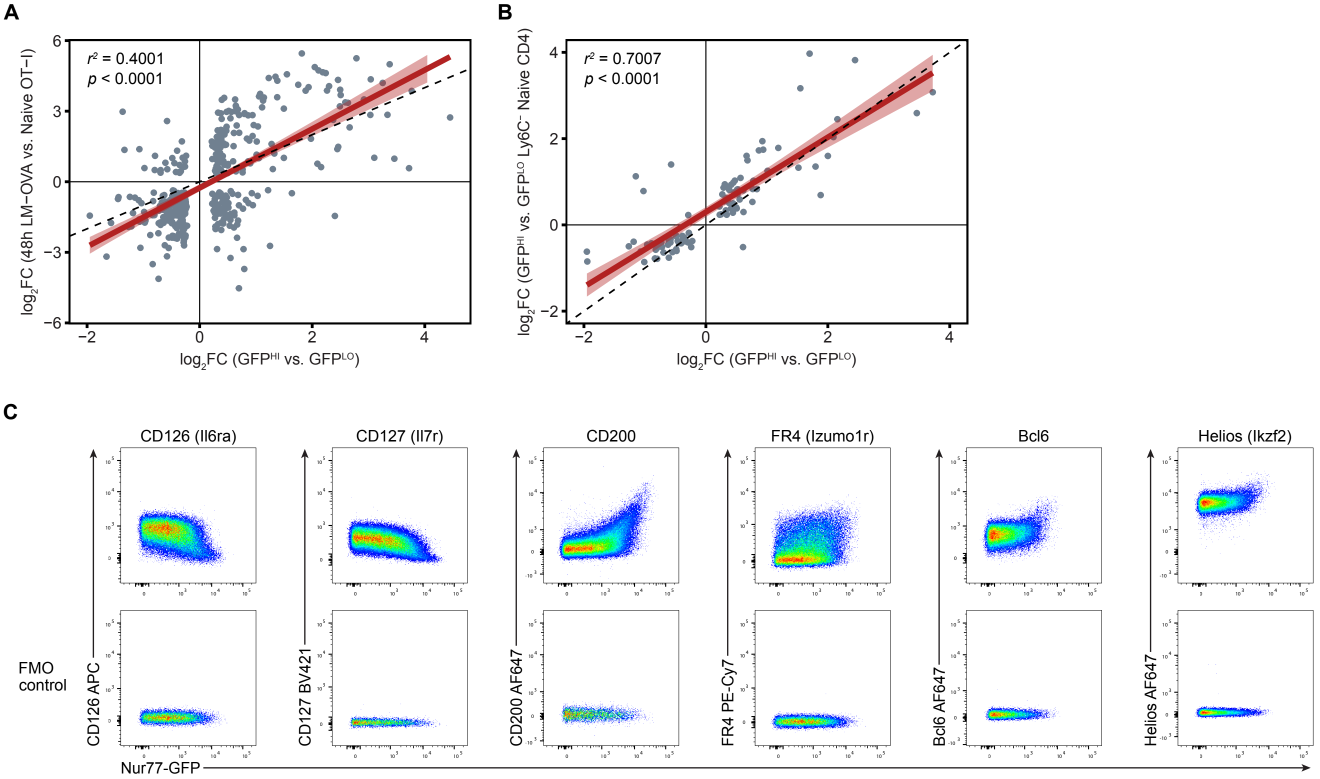

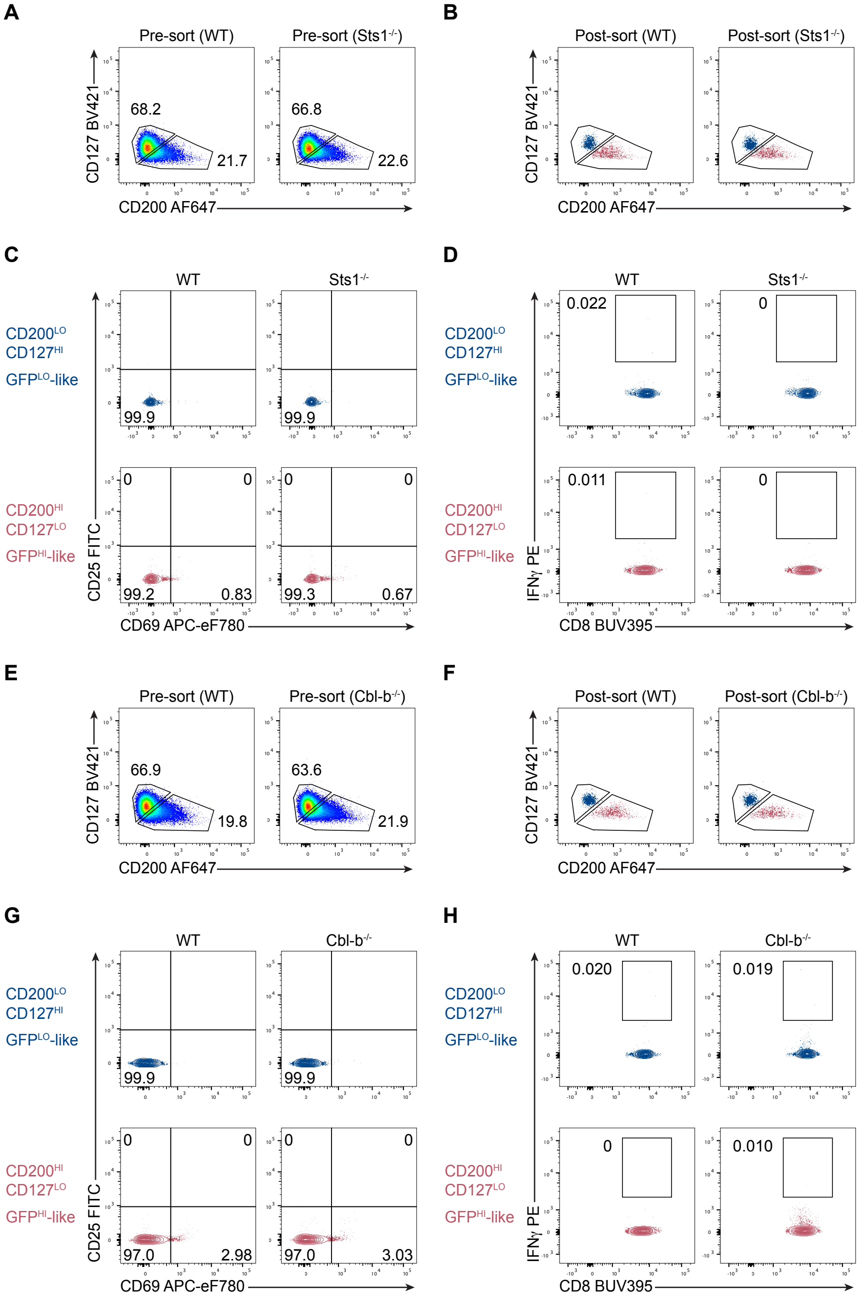
